# Enhancing Polysaccharide Immunity through Intranasal Immunizations

**DOI:** 10.1101/2022.04.19.488813

**Authors:** Kevin Trabbic, Shyamala Ganesan, Ali Fattom, Andrew Lees, Vira Bitko

## Abstract

**Objective:** A model conjugate vaccine incorporating a pneumococcal polysaccharide with EcoCRM_197_ was used to determine vaccine efficacy when administered with our clinical-stage mucosal stimulating adjuvant, NE01. This was the first attempt at combining NE01 with a polysaccharide conjugate vaccine to elicit mucosal immunity against respiratory bacterial infection through intranasal immunizations.

**Results:** Intranasal immunizations using the NE01 adjuvant incorporating a polysaccharide conjugate resulted in the generation of a robust IgG and IgA responses in both serum and mucosa. Our data also demonstrated the active homing of immunological memory cells to the lower respiratory tract as evidenced by an increase of IgG- and IgA-producing memory B-cells in the lungs. Further, intranasal immunization enhanced the induction of a balanced Th1/Th2/ Th17 immune response with clear homing of memory T-cells to the lungs. Serum antibodies generated by this formulation and route of administration demonstrated excellent efficacy in killing *S. pneumonia* in an *in vitro* OPK assay, a biomarker of vaccine efficacy. Our data suggest that further optimization of the dose and schedule as well as evaluation of the efficacy of this new formulation in animal models of colonization and infection are warranted.

## Introduction

Polysaccharide conjugate vaccines administered intramuscularly are able to produce systemic immunity that demonstrates excellent efficacy against invasive disease caused by *Streptococcus pneumonia*. It was also observed that nasal colonization in infants was reduced for a period of 1-2 years post vaccination with the polyvalent conjugate vaccine (Prevnar). However, at the age of 2 years, nasopharyngeal colonization rates were almost identical between children who received the Prevnar vaccine and those who did not. The observed transient protection against colonization following IM vaccination could be attributed to enhanced IgG titers seeping from the sera to the mucosa which could prevent initiation of colonization or suppress existing colonizing bacteria. When the levels of serum IgG wane down, the IgG disappears from the nasopharyngeal mucosa resulting in recolonization. It is suspected that natural colonization aids mucosal immunity by boosting IgG and IgA antibodies to the capsular polysaccharide and cell wall proteins.^1^ Mass immunizations campaigns using approved pneumococcal vaccines have diminished invasive pneumococcal infections. Unfortunately, these IM vaccines had a marginal efficacy against pneumonia and Acute Otitis Media. A new formulation allowing intranasal administration of these vaccines to generate strong mucosal immunity that prevents bacterial colonization and mucosal infections is a major unmet medical need.

Currently, there are no FDA-approved bacterial-based polysaccharide conjugate vaccines that are delivered intranasally. There have been limited attempts focused on generating mucosal immunity using polysaccharide conjugate vaccines through local intranasal immunizations. In one such effort, a polysaccharide conjugate derived from Streptococus pneumoniae used RhinoVax®, a mucosal adjuvant consisting of capylic-capric glycerides.^2^ This immunization approach conferred protection and prolonged survival in mice against a challenge but failed to demonstrate mucosal protection or an equivalent immune response compared to non-adjuvanted intramuscular immunizations. A successful mucosal adjuvant would not only enhance polysaccharide-specific mucosal immunity but would also prevent colonization.

Our NanoVax platform is a clinical-stage adjuvant consisting of NE01, an oil-in-water adjuvant used to induce mucosal and systemic immunity through intranasal delivery. The use of our platform has demonstrated protection and involved the incorporation of protein-based antigens from SARS-CoV-2 (unpublished), HSV, RSV, *Mycobacterium tuberculosis*, anthrax and flu.^3–6^ Additionally, our adjuvant platform contributed to eliminating HSV-2 infection in guinea pigs through a strong cell-mediated and mucosal immunity. In this note, we aim to show the feasibility of incorporating a pneumococcal polysaccharide (PnPs) conjugated to CRM_197_ to enhance polysaccharide specific mucosal immunity.

## Materials and Methods

### Conjugation of Pn18c to CRM197

Pneumococcal capsular polysaccharide 18C was a gift of Inventprise, LLC (Redmond, WA). The polysaccharide was aminated using CDAP activation followed by derivatization with hexanediamine^19^. CRM_197_ (EcoCRM®, Fina Biosolutions) was conjugated to the aminated polysaccharide using thiol-ether chemistry^20^.

### Vaccine Preparation and Mouse Immunizations

The vaccine was prepared by mixing of Pn18C-CRM197 containing 4.0 μg of total polysaccharide with 60% NE01. Mouse immunizations were performed at IBT Bioservices (Rockville, MD) under approved IACUC animal study protocol. Two groups of 7 female CD-1 mice (six to eight week old) were either intranasally immunized with 12 uL of Pn18c-CRM197/NE01 on day 0, 21, 42 or were not immunized (control). On day 70, the following samples were obtained; blood, bronchoalveolar lavage (BAL) fluid, spleens, and lungs.

### Determination of Serum and BAL Pn18C specific IgG and IgA by ELISA

96-well Immulon 4HBX plates were coated with either 1μg/mL of Pn18C or CRM197, blocked with 5% BSA, and serum or BAL samples were added to plates followed by two-fold serial dilutions. Antibodies were detected using either Anti-Mouse IgG-HRP (Jackson Immunoresearch 515-035-071) or Rabbit Anti-Mouse IgA-HRP(Rockland 610-4306). The endpoint titer (EPT) was determined by extrapolating the OD values from the dilution points spanning the cutoff value (3 times the mean of background) and calculating the average.

### Lung and Spleen Cytokine Assay and B-cell ELISPOT

Individual single cells suspensions were prepared by digesting the tissues with collagenase and DNAse, followed by removal of contaminating red blood cells by using 0.8% ammonium chloride with EDTA, washed with media, and resuspended and plated on a 96 well plate at 5×10^5^ cells per well. Cells were stimulated with CRM197 or Pn18c for 72 hr and supernatants were collected and Luminex assays were performed according to manufacturer protocol.

For ELISpot assay, cells suspensions were stimulated with IL-2 (0.5μg/mL) and RD848 (1μg/mL) for 3 days followed by washing and plating onto PVDF ELISpot filter plates coated with anti-mouse IgG or IgA. The cells were incubated for 24 hrs at 37°C, followed by the addition of biotinylated Pn18C. Anti-IgG or Anti-IgA antigen specific cells were detected using streptavidin-HRP. The spots were quantified in AID ELISpot reader.

### Opsonization Killing Assay (OPKA)

The OPKA was performed as previously described.^21^ Briefly, 4.5×10 HL-60 cells/mL were differentiated for 5 days using 0.8% dimethylformamide (DMF) in 10%FBS RPMI-1640 media and adjusted to a final concentration of 1.0×10^7^ cells/mL after washing. Heat-inactivated serum was serially diluted on a 96 well plate and incubated with 10 μL of 5×10^4^ CFU of S. pneumoniae strain OREP18 and incubated for 30 min at room temperature. 10μL of baby rabbit complement and 40 μL of differentiated HL-60 cells were incubated for 45 min at 37°C 5%CO_2_. The assay was stopped by placing the plate on ice for 20 min. 5μL from each well was spotted on a Todd-Hewitt Broth, Yeast (THBY) agar plate followed by an THBY overlay containing triphenyl tetrazolium chloride (TTC). The plates were incubated overnight at 37°C 5%CO_2_ incubator. The colonies were counted and opsonization titers were determined from the interpolation of the serum dilution spanning 50 % killing cutoff determined from S. *pneumoniae* subjected to assay conditions without heat inactivated serum.

## Results

The pneumococcal conjugate used in these immunizations was modeled after an individual component from Prevnar 13, where the capsular polysaccharide Pn18c from *Streptococcus pneumonia* strain OREP18c was conjugated to CRM_197_. The Intranasal immunizations with Pn18c-CRM197/NE01 resulted in a 1.5-fold increase of polysaccharide specific IgG antibodies in serum and BAL in comparison to the specificity to CRM_197_. In the bronchoalveolar fluid (BALf) there was a 2-fold increase in the IgA polysaccharide (PS) specific antibodies compared to the carrier recognition (**Figure 1a-c**). A negligible IgA PnPS (Pn18C-polysaccharide)-specific response was observed in the serum but there was a presence of IgA and IgG antibodies in the upper respiratory tract. Presence of the Pn18c-specific antibodies in the mucosa is encouraging because it provides an evidence of activation of the mucosal immune response, which is not observed with traditional IM polysaccharide conjugate vaccines.

**Figure 1.**
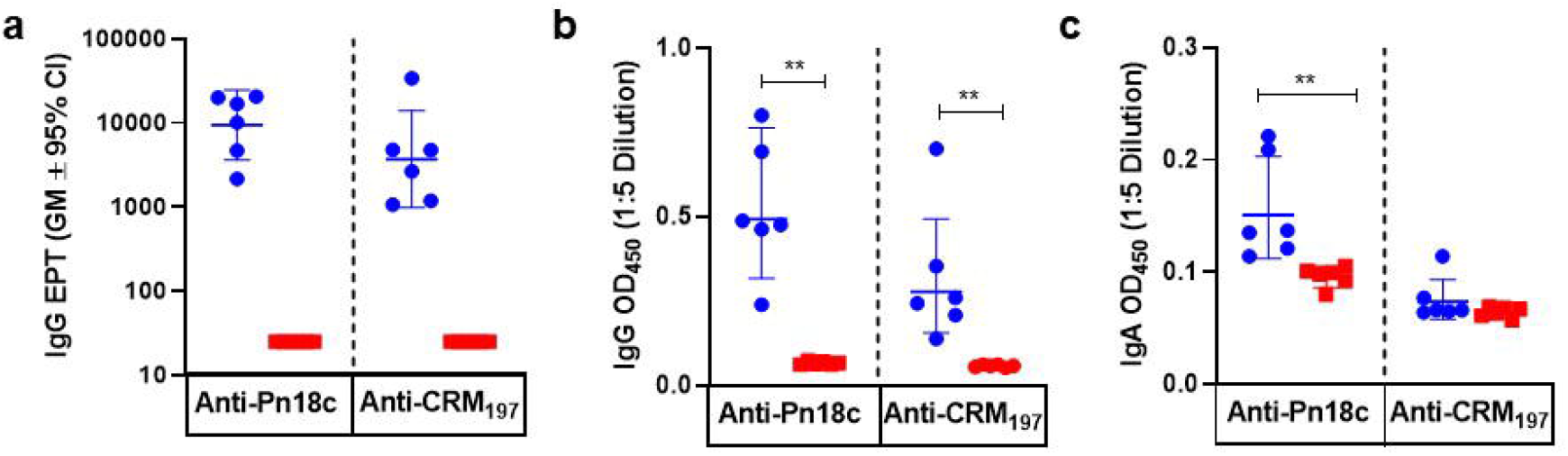
Levels of Pnl8c-specific IgG and IgA after IN vaccination with Pnl8c-CRMI97/NE01. Humoral immune responses elicited to Pn18c or CRM_197_ from IN immunizations of mice. All immunized animals received three immunizations with Pnl8c-CRM_197_/NE01 (blue circles) or PBS as a control (red squares). Anti-sera was serially 2-fold diluted and titers were calculated from the extrapolation of the curve crossing the cut off value, a) Serum IgG EPT between Pn18c compared to CRM_197_ b) BAL IgG OD values c) BAL IgA OD values. Statistical analysis was calculated using Maim-Whitney nonparametric test for unpaired data, \**p*<0. 05, \*\**p*<0.01.

Cellular immunity generated from Pn18c-CRM_197_/NE01 immunizations resulted in a balanced Th1/Th2/Th17 activation, with a significant stimulation of T cells in the lung (**Figure 2a-d**) and spleen (data not shown) by CRM_197_. It is well known that polysaccharides are T cell independent antigens and activation of T cells is largely dependent on antigenic glycopeptides or peptide fragments from CRM_197_ conjugates presented by MHCII. The cytokines released by lung cells upon Pn18c stimulation demonstrated 8-fold increase in IFNy, a 5-fold increase in IL-17, and a 2.2-fold increase in IL-5 compared to control animals. There was greater T cell activation in the lung compared to the spleen (not shown), which suggests Pn18c-CRM_197_ conjugates influence local cellular immunity. Although not statistically significant, the increased levels of IL-17 stimulated by Pn18c could provide support for a superantigen T-cell activation mechanism, through non-specific MHCII interactions.^7^

**Figure 2.**
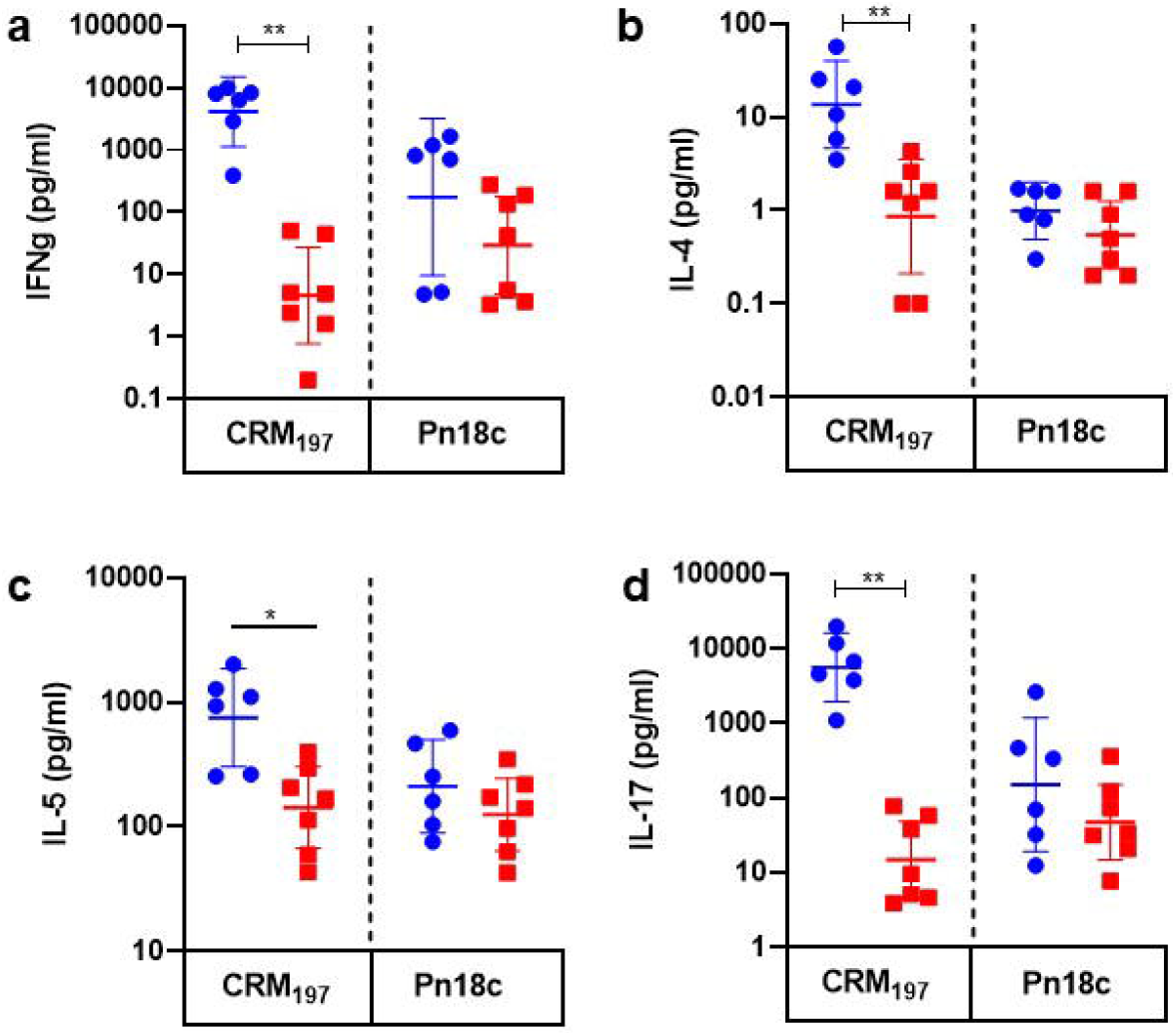
Cytokine quantification in the lungs of Pnl8c-CRM_197_/NE01 IN immunized mice. Comparison of T-cell associated cytokine release between mice immunized with 3-doses of IN Pnl8c-CRM_197_/NE01 (blue circles) or control (red squares). Release of (a) INFγ, (b) IL-4, (c) IL-5, and (d) IL-17 from Pn18c and CRM_197_ stimulated single cell suspension of lungs. Data are presented as the geometric mean with a 95% confidence interval and statistical analysis was calculated between groups using Maim-Whitney nonparametric test for unpaired data, \**p*<0.05, \*\**p*<0. 01.

Further supporting evidence for polysaccharide mucosal and systemic immunity, stimulated B cells isolated from the lung and spleen demonstrated a 3-fold increase of IgG and IgA secreting B cells. An increase in the number of IgG polysaccharide-specific tissue-resident memory B cells suggests there was a skewed IgG class switching evidenced by increased serum and BAL IgG (**Figure 3a**). Additionally, serum from Pn18c-CRM_197_/NE01 intranasally vaccinated animals showed that this vaccination resulted in production of antibodies with the opsonophagocytic killing activity towards *S. pneumoniae* strain OREP18c (**Figure 3b**). These antibodies are a strong indicator that intranasal immunization would produce enhanced immune recognition towards the capsular polysaccharide resulting in protection.

**Figure 3.**
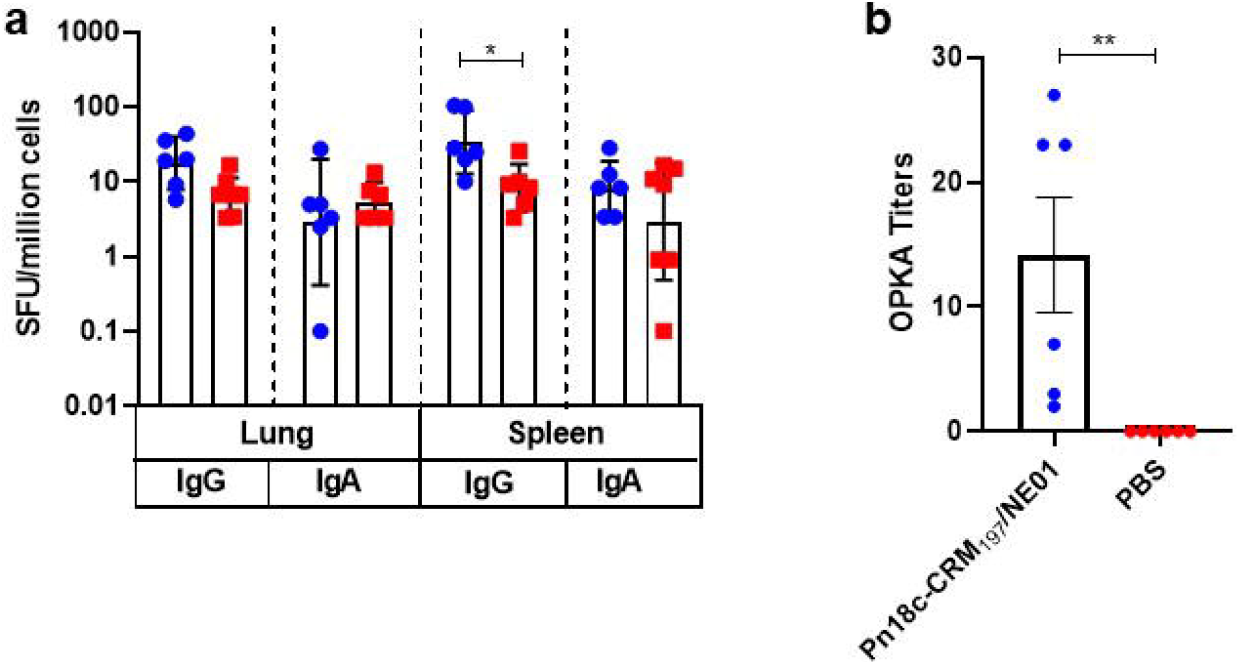
ELISPOT quantification of IgG- and IgA-producing B-cells and OPKA titers. (a) Comparison between the quantification of SFU between animals immunized with 3 doses of IN Pnl8c-CRM_197_/NE01 from cells in the lungs and spleens. Poly-clonal stimulation of B cells were initiated by IL-2 and RD848, followed by Pn18c specific capture of IgG and IgA. (b) OPKA titers using serum from animals immunized with 3 doses of IN Pnl8c-CRM_197_ /NE01 (blue circles) or control (red squares) with Streptococcus *pneumonia* strain OREPl8c. Data are presented as the geometric mean with a 95% confidence interval and statistical analysis was calculated between groups using Mann-Whitney nonparametric test for unpaired data, \**p*<0.05, **p<0.01.

## Discussion

Bacterial polysaccharide conjugate vaccines such as Prevnar or VAXNEUVANCE have been very successful in providing long-term systemic immunity against pneumococcal diseases. These commercial vaccines are administered via intramuscular route with aluminum salts as adjuvants. However, adsorption of protein and/or polysaccharide antigens to elemental aluminum can catalyze degradative processes such as oxidation, hydrolysis, or deamination. ^8, 9^ These degradative processes have been shown to reduce overall immunogenicity generated against polysaccharides by not preserving native structure and structural confirmations. In addition, aluminum-based adjuvants do not induce mucosal immunity and the mucosal effect is considered transitional. The mechanism of action of aluminum-based adjuvants is in stark contrast to our NE01 mucosal adjuvant, where local immunity is stimulated through intranasal immunizations and delivered in an inert chemical environment.

The Pn18c-CRM_197_/NE01 vaccine generated strong humoral immunity with antibodies responses that were strongly skewed towards IgG, which had enhanced selectivity towards Pn18c over CRM_197_. The lack of a robust serum IgA response can be attributed to carrier protein suppression by activating strong Th2 responses, which in turn influence the specific antibody isotype observed. ^10^ The lack of IgA antibodies are not surprising as the production of IgA antibodies from polysaccharide conjugates have shown a strong reduction in IgA detectable in the salivia.^11^ However, based on our previous data using NE01, it has bolstered strong mucosal immunity; homed IgA- and IgG-producing B-cells and Th1/Th2/ Th17 T-cells to mucosa, resulting in full protection against HSV2, RSV, and flu infections with completely or significantly reduced viral titers remaining in the nasal cavitiy.

A fundamental difference between polysaccharide conjugates vaccines and polysaccharide vaccines is the induction of a strong Th2 immunity from the combination of carrier proteins and adjuvant through intramuscular immunizations. After immunizations with Prevnar 13, there is a sustained production of PS-specific IgG responses, however levels of IgA after an initial rapid onset rapidly decline.^1^ Also, bolstered IgA production is not observed by boosting with Pneumovax 23 suggesting the induction of Th2 immunity favoring additional differentiation of IgG producing B cells over IgA. ^12^ The traditional mechanism of polysaccharide conjugates involves only the peptide portion of the carrier protein to be presented to T cells via MHC II and, activated T cells influence antibody secreting cells. However, an alternative mechanism has been demonstrated where PS is processed creating glycopeptides that are able to activate carbohydrate-specific helper CD4+ T cell clones known as Tcarbs.^13^ While the T cell activation was predominately influenced by peptide stimulation from the carrier protein (Figure 2) processed glycopeptide portions were likely presented to T cells evidenced by the direct stimulation from Pn18c alone. Our results revealed there was a balanced Th1/Th2/Th17 that was detectable in the lungs. These results are supported by literature as lung Th17 cells play a key role in protection against pneumococcal infections.^1,14^ IL-17 generation from vaccines have been understudied however, IL-17 has been shown to be associated with the recruitment of neutrophils to subsequently eradicate pathogens. It is also, in association with IL-22 boosts the expression of β-defensin 2, which faciliates IgA transport into mucosa by promoting pIgR expression which in turn promotes elimination of pathogens at mucosal surfaces preventing pathogenic nasal colonization. ^15,16,17^. Additionally, there is a correlation between reduced IL-17a levels in children and a higher density of nashpharyngeal carriage,^1,18^ which highlights the importance of mucosal immunity and the possibility of its activation through intranasal immunization with polysaccharide nanoemulsion-adjuvanted vaccine.

Locally primed immunity through our adjuvant system is critical in developing a robust local mucosal immune response and memory. The local immunity can confer protection for the short term while the local mucosal tissue-resident memory cells will be activated immediately after exposure to the pathogen and act as the first line of defense against the invading pathogen. The motivation of this study was to determine the feasibility of our intranasal adjuvant platform using a model polysaccharide conjugate by enhancing mucosal and systemic immune response to polysaccharide antigens. Additional studies to explore dosing, regimen, protection in appropriate animal models, as well as comaparing to IM immunized animals are warranted. Our adjuvant system has shown to be a prime candidate for polysaccharide conjugates, especially polysaccharides that contain liable linkages.

## Limitations

The study demonstrated the induction of mucosal and systemic immune response by intranasl delivery of polysaccharide antigens. However, there was no direct comparison of intramuscular route delivery of the same polysaccharide antigen in this experiment. Additionally, in direct efficacy read out using OPK assay is demonstrated rather than using appropriate animal challenge models.

## Declarations

### Ethics Approval and Consent to Participate

Not applicable

### Consent for Publication

Not applicable

### Availability of Data and Materials

All data generated or analyzed during this study are available from the corresponding author on reasonable request.

### Author’s Contributions

KT performed the in vitro testing, analyzed data and wrote the manuscript draft; SG designed the study, prepared vaccine formulations and reviewed the manuscript; AF and VB designed and supervised the study and revised the manuscript. AL provided the polysaccharide and EcoCRM and reviewed the manuscript. All authors read the manuscript, revised and accepted it.

## Acknowledgments

We are thankful to Hugo Acosta, Kallista Orange, Chris Brigolin and Maggie Lugin for processing the single cell suspensions and performing the assays to determine the cell-mediated and systemic immune responses.

## References

1. Clarke ET, Williams NA, Dull PM, Findlow J, Borrow R, Finn A, et al. Polysaccharide-protein conjugate vaccination induces antibody production but not sustained B-cell memory in the human nasopharyngeal mucosa. Mucosal Immunol 2013; 6:288–96.

2. Jakobsen H, Saeland E, Gizurarson S, Schulz D, Jonsdottir I. Intranasal immunization with pneumococcal polysaccharide conjugate vaccines protects mice against invasive pneumococcal infections. Infect Immun 1999; 67:4128–33.

3. Stanberry LR, Simon JK, Johnson C, Robinson PL, Morry J, Flack MR, et al. Safety and immunogenicity of a novel nanoemulsion mucosal adjuvant W805EC combined with approved seasonal influenza antigens. Vaccine 2012; 30:307–16.

4. Bernstein DI, Cardin RD, Bravo FJ, Hamouda T, Pullum DA, Cohen G, et al. Intranasal nanoemulsion-adjuvanted HSV-2 subunit vaccine is effective as a prophylactic and therapeutic vaccine using the guinea pig model of genital herpes. Vaccine 2019; 37:6470–7.

5. O’Konek JJ, Makidon PE, Landers JJ, Cao Z, Malinczak CA, Pannu J, et al. Intranasal nanoemulsion-based inactivated respiratory syncytial virus vaccines protect against viral challenge in cotton rats. Hum Vaccin Immunother 2015; 11:2904–12.

6. Ahmed M, Smith DM, Hamouda T, Rangel-Moreno J, Fattom A, Khader SA. A novel nanoemulsion vaccine induces mucosal Interleukin-17 responses and confers protection upon Mycobacterium tuberculosis challenge in mice. Vaccine 2017; 35:4983–9.

7. Torres BA, Perrin GQ, Mujtaba MG, Subramaniam PS, Anderson AK, Johnson HM. Superantigen enhancement of specific immunity: antibody production and signaling pathways. J Immunol 2002; 169:2907–14.

8. Wittayanukulluk A, Jiang D, Regnier FE, Hem SL. Effect of microenvironment pH of aluminum hydroxide adjuvant on the chemical stability of adsorbed antigen. Vaccine 2004; 22:1172–6.

9. HogenEsch H, O’Hagan DT, Fox CB. Optimizing the utilization of aluminum adjuvants in vaccines: you might just get what you want. NPJ Vaccines 2018; 3:51.

10. Avci FY, Li X, Tsuji M, Kasper DL. A mechanism for glycoconjugate vaccine activation of the adaptive immune system and its implications for vaccine design. Nat Med 2011; 17:1602–9.

11. Zhang Q, Choo S, Everard J, Jennings R, Finn A. Mucosal immune responses to meningococcal group C conjugate and group A and C polysaccharide vaccines in adolescents. Infect Immun 2000; 68:2692–7.

12. Crowther RR, Collins CM, Conley C, Lopez OJ. Rapid kinetics of serum IgA after vaccination with Prevnar((R))13 followed by Pneumovax((R))23. Heliyon 2017; 3:e00255.

13. Sun X, Stefanetti G, Berti F, Kasper DL. Polysaccharide structure dictates mechanism of adaptive immune response to glycoconjugate vaccines. Proc Natl Acad Sci U S A 2019; 116:193–8.

14. Wang Y, Jiang B, Guo Y, Li W, Tian Y, Sonnenberg GF, et al. Cross-protective mucosal immunity mediated by memory Th17 cells against Streptococcus pneumoniae lung infection. Mucosal Immunol 2017; 10:250–9.

15. Jaffar Z, Ferrini ME, Herritt LA, Roberts K. Cutting edge: lung mucosal Th17-mediated responses induce polymeric Ig receptor expression by the airway epithelium and elevate secretory IgA levels. J Immunol 2009; 182:4507–11.

16. Ye P, Rodriguez FH, Kanaly S, Stocking KL, Schurr J, Schwarzenberger P, Oliver P, Huang W, Zhang P, Zhang J, Shellito JE, Bagby GJ, Nelson S, Charrier K, Peschon JJ, Kolls JK. Requirement of interleukin 17 receptor signaling for lung CXC chemokine and granulocyte colony-stimulating factor expression, neutrophil recruitment, and host defense. J Exp Med. 2001;194:519–527.

17. Liang SC, Tan XY, Luxenberg DP, Karim R, Dunussi-Joannopoulos K, Collins M, Fouser LA. Interleukin (IL)-22 and IL-17 are coexpressed by Th17 cells and cooperatively enhance expression of antimicrobial peptides. J Exp Med. 2006;203:2271–2279

18. Hoe E, Boelsen LK, Toh ZQ, Sun GW, Koo GC, Balloch A, et al. Reduced IL-17A Secretion Is Associated with High Levels of Pneumococcal Nasopharyngeal Carriage in Fijian Children. PLoS One 2015; 10:e0129199.

19. Lees A, Barr JF, Gebretnsae S. Activation of Soluble Polysaccharides with 1-Cyano-4-Dimethylaminopyridine Tetrafluoroborate (CDAP) for Use in Protein-Polysaccharide Conjugate Vaccines and Immunological Reagents. III Optimization of CDAP Activation. Vaccines (Basel). 2020 Dec 18;8(4):777

20. Lees A, Nelson BL, Mond JJ. Activation of soluble polysaccharides with 1-cyano-4-dimethylaminopyridinium tetrafluoroborate for use in protein-polysaccharide conjugate vaccines and immunological reagents. Vaccine. 1996 Feb;14(3):190–8.

21. Burton RL, Nahm MH. Development and validation of a fourfold multiplexed opsonization assay (MOPA4) for pneumococcal antibodies. Clin Vaccine Immunol 2006; 13:1004–9.

